# A metabolic shift to glycolysis promotes zebrafish tail regeneration through TGF–β dependent dedifferentiation of notochord cells to form the blastema

**DOI:** 10.1101/2020.03.03.975318

**Authors:** Jason W. Sinclair, David R. Hoying, Erica Bresciani, Damian Dalle Nogare, Carli D. Needle, Weiwei Wu, Kevin Bishop, Abdel G. Elkahloun, Ajay Chitnis, Paul Liu, Shawn M. Burgess

## Abstract

Mammals are generally poor at tissue regeneration, in contrast, fish maintain a high capacity for regenerating complex tissues after injury. Using larval zebrafish, we show that tail amputation triggers an metabolic shift to glycolysis in cells surrounding the notochord as they reposition to the amputation site. Blocking glycolysis prevents the fin from regenerating after amputation due to the failure to form a normal, pluripotent blastema. We performed a time series of scRNA-sequencing on regenerating tails under normal conditions or in the absence of glycolysis. Strikingly, we detected a transient cell population in the single cell analysis that represents notochord sheath cells undergoing a TGF–β dependent dedifferentiation and epithelium-to-mesenchyme transition to become pluripotent blastema cells. We further demonstrated that the metabolic switch to glycolysis is required for TGF–β signaling and blocking either glycolysis or TGF–β receptors results in aberrant blastema formation through the suppression of essential EMT mediators such as *snai1*.

## Introduction

Many non-mammalian vertebrates have the ability to regenerate complex appendages following loss or injury. Appendage regeneration is a process involving the coordinated interaction of multiple cell types following injury: there is an immune response and the establishment of a wound epithelium or a specialized epithelial structure known as the apical epidermal cap (AEC), and then signaling that emanates from the newly formed wound epithelium / AEC through growth factors such as TGF–β and FGF. These early responses promote cell migration and proliferation, and are required for blastema formation, the key step leading to complete tissue restoration of an appendage. The blastema, which has been described as mesenchymal in nature, is a mass of multipotent cells capable of proliferating and differentiating to replace lost tissue. Although mammals typically have a very limited capacity to form blastemas and thus cannot regenerate appendages, there are a few examples where blastema formation can occur in mammals. Two species of African spiny mouse have been reported to form blastema-like structures during skin shedding [1] and digit tip regeneration in mice also involves the formation of a blastema from fate-restricted progenitors [2]. Human children are also capable of digit tip regeneration, suggesting limited blastema formation is at least possible in humans [3]. Interestingly, in contrast to most commonly studied mouse strains, the MRL mouse does form a blastema around ear punctures, suggesting that tissue regeneration in mammals could be enhanced or even initiated with a better understanding of the pathways underlying blastema formation [4]. Although progress has been made in determining cellular signals required for blastema formation and maintenance, the mechanisms by which apparently differentiated cells dedifferentiate to become the blastemal mesenchyme remains poorly understood.

A frequently observed phenomenon during regeneration is metabolic switching, or the preferential utilization of glycolysis as an ATP source uncoupled from oxidative phosphorylation. A shift towards glycolytic metabolism has been proposed as a mediator of early tail regeneration in both xenopus and lizards [5, 6] and has been shown to be essential in zebrafish heart regeneration [7]. Beyond regeneration, there are many other examples of metabolic switching regulating cellular function. For example, human embryonic stem cells rely on glycolysis in a mildly hypoxic environment for self-renewal. Upon exposure to oxygen, these cells increase oxidative phosphorylation and quickly lose pluripotency [8]. Inflammatory macrophages rely primarily on glycolysis while repair macrophages utilize oxidative phosphorylation, and switching in these cells activates distinct transcriptional profiles [9]. Highly proliferative cells, including many types of cancer, utilize aerobic glycolysis, termed the Warburg effect, as their main energy source [6]. In addition to increased glucose uptake, cancer cells utilizing aerobic glycolysis often have increased ROS production, which has also been shown to be a mediator of tissue regeneration [10-14]. Proposed reasons for the Warburg effect include faster, albeit less efficient ATP production, adaption to hypoxic conditions, shunting of glucose metabolites to anabolic pathways, manipulation of the microenvironment, and metabolites used for cell signaling [15]. With respect to cell signaling, it is becoming increasingly clear that metabolic reprogramming plays a role in promoting an epithelial to mesenchyme transition (EMT) in tumor cells [16]. EMT is the process by which epithelial cells lose polarity, dismantle cell-cell junctions, and gain migratory properties to become mesenchymal stem cells. EMT is a crucial process in normal development, wound healing or tissue regeneration, and cancer progression. Several signaling pathways, including TFG−β, Notch, and Wnt can induce EMT through the transcription factors Snail, Zeb, and Twist [17]. Despite aerobic glycolysis being observed in multiple biological contexts, the specific purpose for uncoupling aerobic oxidative phosphorylation and its relationship to EMT have not been established.

It has been shown that the zebrafish *Dario rerio* can regenerate many tissues after injury or amputation including the fins [18], the heart [19], and the spinal cord [20]. In zebrafish embryos, tail amputation through the posterior trunk of the animal results in repositioning of notochord cells to the amputation site, followed by blastema formation and regeneration of several different tissues including the notochord, the neural tube, blood vessels, and the caudal fin fold [21]. Using this tail excision model, previous reports have shown that sonic hedgehog and EGFR signaling are required for regeneration [21, 22]. However, the full genetic pathway for blastema formation following repositioning of notochord cells has not been determined. Here, we show that a metabolic switch from oxidative phosphorylation to aerobic glycolysis is obligate for fin regeneration. Blocking glycolysis has no gross effects on early larval development under normal conditions, but results in a complete failure of the caudal fin to regenerate following amputation. During regeneration, notochord cells specifically increase glucose uptake and display altered mitochondrial morphology and oxidation state as they are repositioned to the notochord bud, indicative of metabolic reprogramming. Subsequently, sheath cells on the periphery of the notochord bud undergo EMT, dedifferentiate, and contribute to blastema formation. Single-cell RNAseq analyses revealed that inhibiting glycolysis resulted in the failed induction of TGF–β ligands and *snai1* expression essential to triggering the EMT during blastema formation. Similarly, direct inhibition of TGF–β resulted in a failure to form the blastema through suppression of the EMT. Together our data indicate that a metabolic shift to glycolysis is a key event regulating normal blastema formation by triggering TGF–β signaling in notochord sheath cells. The metabolic shift towards glycolysis and EMT that takes place in the notochord bud suggests potential parallels between tissue regeneration and tumor biology, including ways to regulate or inhibit them to our advantage.

## Materials and Methods

### Zebrafish Husbandry

Adult zebrafish were maintained at 28?°C with a 14 h light / 10 h dark cycle. Embryos collected from crosses were staged as previously described [23]. All animal experiments were performed in compliance with NIH guidelines for animal handling and research (Protocol G-01-3).

### Microscopy

Embryos were anesthetized with MS-222 (Sigma) and embedded in 0.8% agarose (Fisher) for imaging. Epifluorescence images were acquired with an Axiovert 200M (Zeiss) and an Orca-ER camera (Hamamatsu). Bright field images were acquired with an MZ16F L3 (Leica) and Axio Cam HR (Zeiss). Confocal images were acquired with an LSM 880 (Zeiss). Images were analyzed in FIJI [24, 25].

### Time-lapse microscopy

For time-lapse imaging, embryos were anesthetized in holtfreter’s buffer (59 mM NaCl, 0.88 mM CaCl_2_·2H_2_0, 0.67 mM KCl, 5 mM HEPES) containing 600µM MS-222 (Sigma) and embedded in 1% low-melt agarose (NuSieve GTG) containing MS-222 on glass-bottomed microwell dishes (MatTek 35mm). Imaging was performed using a Nikon W1 spinning disc confocal microscope with a 20X 0.75NA dry objective lens. After acquisition, image sequences were bleach corrected using histogram normalization [26], and channels merged in FIJI.

### Plasmid synthesis

Plasmids were generated with multisite gateway technology (Invitrogen). To obtain Tg(*actb2*:mito-GFP), we generated a middle entry clone with the human COXVIII mitochondrial localization sequence (MLS) fused upstream of GFP [27], which we recombined with p5E−βactin2, p3E-polyA, and pDestTol2pA2 from the tol2 kit [28]. To generate Tg(*actb2*:mito-roGFP), a middle entry clone containing the zebrafish COXVIII MLS and a 3’ entry clone containing the roGFP sequence were recombined with p5E−βactin2 and pDestTol2pA2 from the tol2 kit.

### Generation of transgenic zebrafish

To synthesize *tol2* mRNA, pCS2FA-transposase was linearized using NotI (New England Biolabs), then purified using a PCR purification kit (Qiagen). mRNA was synthesized with mMessage mMachine SP6 Kit (Ambion) and purified with RNA clean and concentrator kit (Zymo Research). 1-cell stage embryos were injected with 50pg *tol2* mRNA and 25 pg plasmid. Injected embryos expressing mosaic GFP were grown to adults and screened for germline transmission.

### Pharmacological studies

To inhibit glycolysis, 2-Deoxy-D-glucose (Sigma) powder was added to holtfreter’s buffer to a final concentration of 100 mM. To inhibit mitochondrial protein import, a 50 mM stock of Mitoblock-6 in DMSO (Calbiochem) was diluted into holtfreter’s buffer and 1% DMS0 to a final concentration of 2.5 µM. To inhibit TGF–β signaling, a 20 mM stock of SB431542 (Sigma) in DMSO was diluted into holtfreter’s solution to a final concentration of 50 µM. For development assays, treatments were initiated at 1 DPF and continued until 4 DPF when embryos were analyzed. For regeneration assays, treatments were initiated 2 hrs prior to amputation and continued throughout the duration of the experiment unless otherwise noted. Inhibitors were washed out and replaced every 24 hrs for all experiments.

### Tail area quantification

Tail amputations were performed as previously described [29]. Relative tail size was quantified using the region of interest method in FIJI. Tails were measured from the posterior end of the notochord to the posterior end of the fin.

### In situ hybridization

In situ hybridization was performed as previously described [30]. Primers used to amplify template cDNA were *snai1a*: For-5’ GCGGGTCCAAATGGACGTAAACC, Rev-5’ TGGCGGCAGTAACTGCAGC; *msx3*: For-5’ AGTGCGCTCTGCTTGGAGAGC, Rev-5’ GTGCTTCAGTCTGTTAAGACAAATAATACATTCCGTAGG. A T7 promoter was included on the reverse primers. Probes were synthesized with T7 polymerase and DIG labeling mix (Roche 10881767001, 11277073910).

### Immunofluorescence

Embryos were fixed in 4% formaldehyde for 45 minutes at room temperature, washed with PBST, and incubated in 1 mg/mL collagenase for 2 hours. Embryos were washed with PBST and frozen in acetone at -20°C for 10 min. Embryos were washed in PBST and incubated in 5% goat serum (Thermofisher NC9270494) in BDP (PBS/0.1% BSA/1%DMSO) for 1h. Embryos were incubated with anti-phospho-SMAD2 primary antibody (Cell Signaling E8F3R) diluted 1:800 in BDP overnight at 4°C, followed by incubation with secondary antibody (Alexa Fluor 488 A-11008) diluted in 1:500 BDP for 1h at RT. Nuclei were stained by incubating with Hoechst 33342 (Thermofisher) diluted 1:2000 in PBST for 20 minutes at RT.

### 2-NBDG uptake

Embryos were incubated in 500 µM 2-NBDG (Cayman) in Holtfreter’s buffer for 2 hours. Embryos were then anesthetized with MS-222 and embedded in 0.8% agarose for microscopy.

### Heat Map generation

Heat maps were generated using Heatmapper (http://www2.heatmapper.ca/).

### Gene List Analyses

Genes list analyses were performed using the Panther Classification System 14.1 (www.pantherdb.org) [31]. Statistical Overrepresentation tests were performed with Fisher’s exact test and corrected with False Discovery Rate.

### Single cell RNA-seq

Tails were amputated at the posterior end of the pigment gap to initiate regeneration. Regenerating tails were collected by performing a second amputation at the anterior end of the pigment gap at 24 or 48 HPA. 80 tails were collected per sample. Amputated tails were dissociated for single cell RNA-seq analysis using a previously described protocol with 1% BSA used in place of FBS [32]. Following dissociation to single cells, cell viability and concentration was analyzed using a Luna cell counter. Gel Bead-in-Emulsions were prepared by loading 10,000 cells per sample in 46.6 µL DMEM + 1% BSA onto a Chromium Chip B (10x Genomics 1000073) and run using the Chromium Controller (10x Genomics). cDNA libraries were generated with Chromium Single Cell 3’ GEM, Library and Gel Bead Kit V3 (10x Genomics). Libraries were sequenced using the NextSeq 500/550 High Output Kit v2.5 (Illumina) on an Illumina NextSeq 550. Reads were aligned to the zebrafish genome and feature-barcode matrices were generated using Cell Ranger (10x Genomics). T-SNE plots and differential gene expression lists were generated with Loupe Viewer (10x Genomics). All data was deposited to GEO under accession GSE145497.

### Statistical analyses

T-tests and ANOVA analyses were performed in Graphpad Prism 8. Experiments were repeated a minimum of three times for statistical analyses. Differential gene expression from scRNAseq data was determined using Loupe Viewer software (10x genomics), which employs a negative binomial exact test and Benjamini-Hochberg correction for multiple tests [33].

## Results

### Tail regeneration following amputation

Deep amputation through the posterior trunk, or the tail, of the larval zebrafish results in regeneration of several types of tissue including the notochord, neural tube, blood vessels, and the caudal fin-fold (Figure 1A). Following amputation, notochord cells migrate to the injury plane and form a small “bud” (Figure 1B), a step necessary to initiate cell proliferation [21]. To gain a better understanding of cell proliferation both spatially and temporally during tail regeneration, we utilized the dual Fucci transgenic zebrafish that expresses cerulean in cells within the S, M, or G2 phases and RFP in cells in G0/1 [34]. We observed proliferating cells in and around the notochord bud 3 hours post-amputation (HPA) (Figure 1C and 1D). Proliferation peaked at 24 HPA and persisted in the bud and other regions of the regenerating tail through 48 HPA. In agreement with previous reports, the blastema is evident by 48 HPA, as shown by strong induction of the blastema/mesenchyme marker *msx3* (Figure 1E). At this timepoint proliferating cells were observed primarily outside of the residual notochord bud, overlapping a region consistent with blastema cells (Figure 1C). This suggests that at least a portion of proliferating cells at 48 HPA are blastemal. By 72 HPA proliferation had significantly decreased and by 96 HPA very few cells in S, M, or G2 phase were detectable. Fin area steadily increased throughout the time-course, irrespective of the number of proliferating cells present, indicating that cell migration, proliferation, differentiation, and hypertrophic cell growth all likely contribute to fin regrowth between 3 and 96 HPA (Figure 1D).

**Figure 1:**
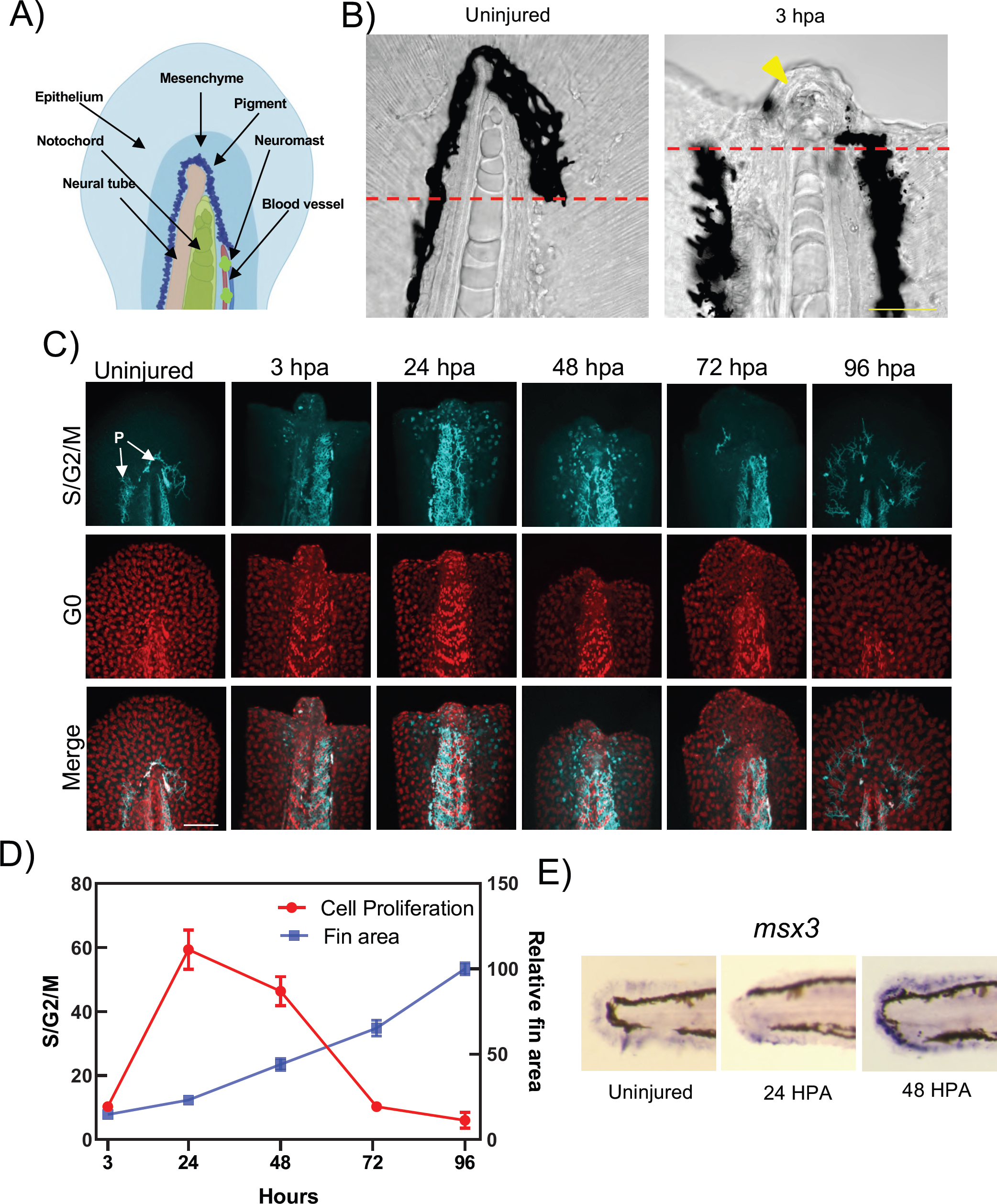
Tail regeneration following amputation in zebrafish embryos. A) An illustration showing tissues that are regenerated following tail amputation. B) Left, a normal tail fin with the amputation plane indicated with a dashed red line. Right, an image depicting the notochord bud formation (yellow arrowhead) at 3 HPA. Scale bar = 50 µM. C). Tail images of the Dual Fucci fish transgenic fish following tail amputation at indicated timepoints in hours post ablation. Cells in S, G2, and M phase are depicted by cyan fluorescence in the nucleus. Cells in G0/1 phase are depicted in red. P indicates autofluorescent pigment cells which are not expressing Tg(*ubb*:Cerulean-gmnn). Scale bar = 100 µM. D) Quantification of cell proliferation and tail area during regeneration. E) In situ hybridization of the blastema marker *msx3* in uninjured animals and at 24 or 48 HPA.

### Aerobic glycolysis is essential for regeneration

Metabolic switching has been shown to play a key role in immune cell responses [35], stem cell maintenance [8, 36], and cancer progression [37]. To understand if it also has a role in regeneration, we impaired either mitochondrial function or glycolysis during development and tail regeneration. We treated 1 day post-fertilization (DPF) embryos with the mitochondrial import inhibitor mitoblock-6 (MB-6), which has been shown to decrease respiration [38]. Treatment resulted in severe developmental abnormalities as previously reported (Supplementary Figure 1A) [39]. We also observed a reduction in caudal fin size, consistent with an essential role for respiration during early development (Supplementary Figure 1B, 1C). In contrast, 2-DG, a well-described glycolysis inhibitor, had no obvious effect on embryo development, and we observed no significant differences in caudal fin size between control and 2-DG treated animals (Figures 2A, 2B and Supplementary Figure 1D). Next, we amputated caudal fins of embryos treated with MB-6. MB-6 treated embryos that survived until 7 DPF had a modest reduction in fin size following amputation but appeared to regenerate mostly normally (Supplementary Figures 1E, 1F). However, in larvae treated with 2-DG, fins completely failed to regenerate compared to control animals (Figure 2C, 2D). Washing out 2-DG prior to 72 HPA resulted in a partial to complete re-initiation of fin regeneration depending on duration of treatment, indicating that the effect of 2-DG treatment was reversible (Figure 2D). Additionally, 2-DG treatment resulted in a decrease of proliferating cells in the regenerating tail (Figure 2E, 2F). Interestingly, adding 2-DG to embryo media 24 HPA resulted in an attenuated phenotype while addition after 48 HPA resulted in no phenotype, indicating that there is an obligate shift to glycolysis early in the regenerative process, prior to blastema formation (Figure 2G) but it is not necessary in later stages of regeneration even though there is still significant cell division taking place (Figure 1C). This suggests that a shift to glycolysis is not purely a function of re-entering the cell cycle. *In situ* hybridization of the blastema marker *msx3* showed that inhibition of glycolysis resulted in the loss of blastema formation, indicating that a metabolic switch to glycolysis is absolutely required for initiating normal blastema development (Figure 2H). Together, these results indicate that while oxidative phosphorylation is essential for normal development and glycolysis is not, there is a obligate shift to glycolysis during the early stages of blastema formation and tail regeneration.

**Figure 2:**
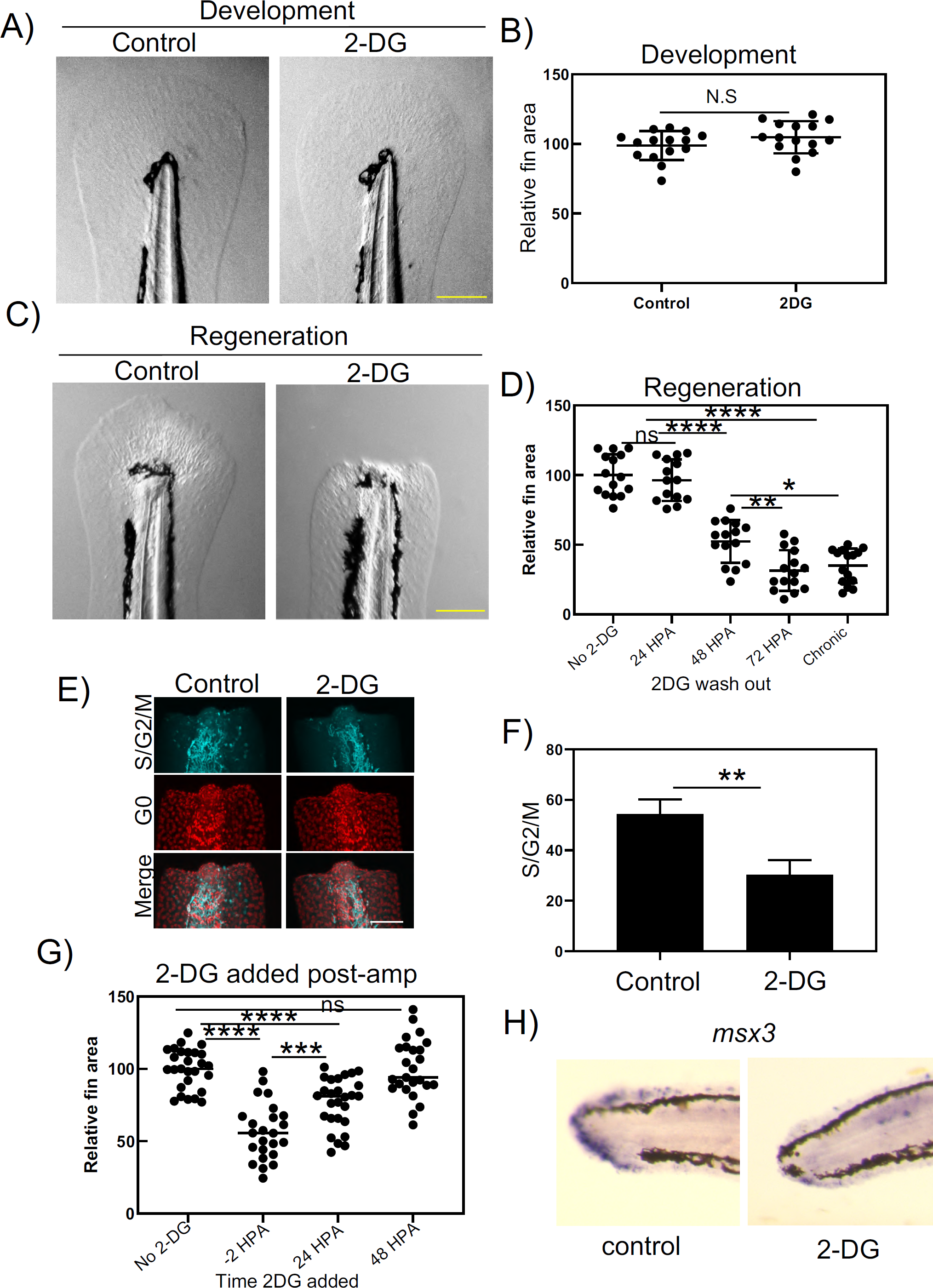
Blocking glycolysis inhibits tail regeneration. A) Images of 4 DPF embryo tails. Embryos were untreated (control) or treated with 2-DG from 1 to 4 DPF. Scale bar = 200 µM. B) Quantification of tail surface area of untreated or 2-DG treated embryos. N.S., not significant (T-test). C) Images of 7 DPF embryo tails 4 days post-amputation. Embryos were untreated (control) or treated with 2-DG from 2 hours before the amputation and throughout the duration of the experiment. Scale bar = 200 µM. D) Quantification of tail surface area of untreated or 2-DG treated embryos 96 HPA. X axis indicates at what timepoint 2-DG was washed out of media. ****P < 0.0001, ***P < 0.001, **P < 0.01, *P < 0.05 (one-way ANOVA). E) Confocal images untreated and 2-DG treated Fucci fish embryo tails at 24 HPA. Scale bar = 100 µM. F) Quantification of cells in S, G2, or M phase in untreated or 2-DG treated fucci embryo tails 24 HPA. **P < 0.01 (T-test). 2-DG significantly repressed cell division. G) Quantification of the tail surface area for untreated or 2-DG treated embryos at 96 HPA. X axis indicates the timepoint that 2-DG was added to the media. ****P < 0.0001, ***P < 0.001 (one-way ANOVA). H) In situ hybridization of *msx3* in untreated and 2-DG treated embryos at 48 HPA.

### Notochord cells increase glucose uptake and display hyperoxidized, fragmented mitochondria following tail amputation

In order to understand which tissues may be undergoing a metabolic switch to glycolysis during caudal fin regeneration, we pulsed embryos with the fluorescent glucose analog 2-(N-(7-Nitrobenz-2-oxa-1,3-diazol-4-yl)Amino)-2-Deoxyglucose (2-NBDG). Within 3 hours post-amputation, we observed a significant increase in the import of 2-NBDG into notochord cells streaming into the notochord bud compared to notochord cells of uninjured animals (Figures 3A, 3B). By 24 HPA glucose uptake was primarily observed in the notochord bud (Figures 3A, 3C). Both cancer cells and stem cells utilizing glycolysis have increased mitochondrial fragmentation compared to cells utilizing oxidative phosphorylation [40], therefore we analyzed mitochondrial morphology by generating a transgenic zebrafish line, Tg(*actb2*:mls-EGFP), which expressed GFP with the Cox VIII mitochondrial localization sequence fused to the N-terminus thereby making the mitochondria fluorescent [27]. Interestingly, we observed that mitochondria began fragmenting as notochord cells entered the bud, again suggesting a glycolytic switch was taking place in these cells (Figure 3D, Supplementary movie 1). In addition to fragmented mitochondria, cancer cells can also show an increase in mitochondrial ROS [41]. To probe mitochondrial ROS production, we generated a second mitochondrial reporter line, Tg(*actb2*:mls-roGFP), expressing a redox sensitive GFP. We observed an increase in the oxidization state of mitochondria, indicative of increased ROS production, specifically within and directly anterior to the notochord bud, similar to the regions of increased glucose uptake and mitochondrial fragmentation (Figure 3E, 3F, and 3G) [42]. Taken together, these data demonstrate that notochord cells undergo a metabolic shift to glycolysis as they reposition toward the amputation plain.

**Figure 3:**
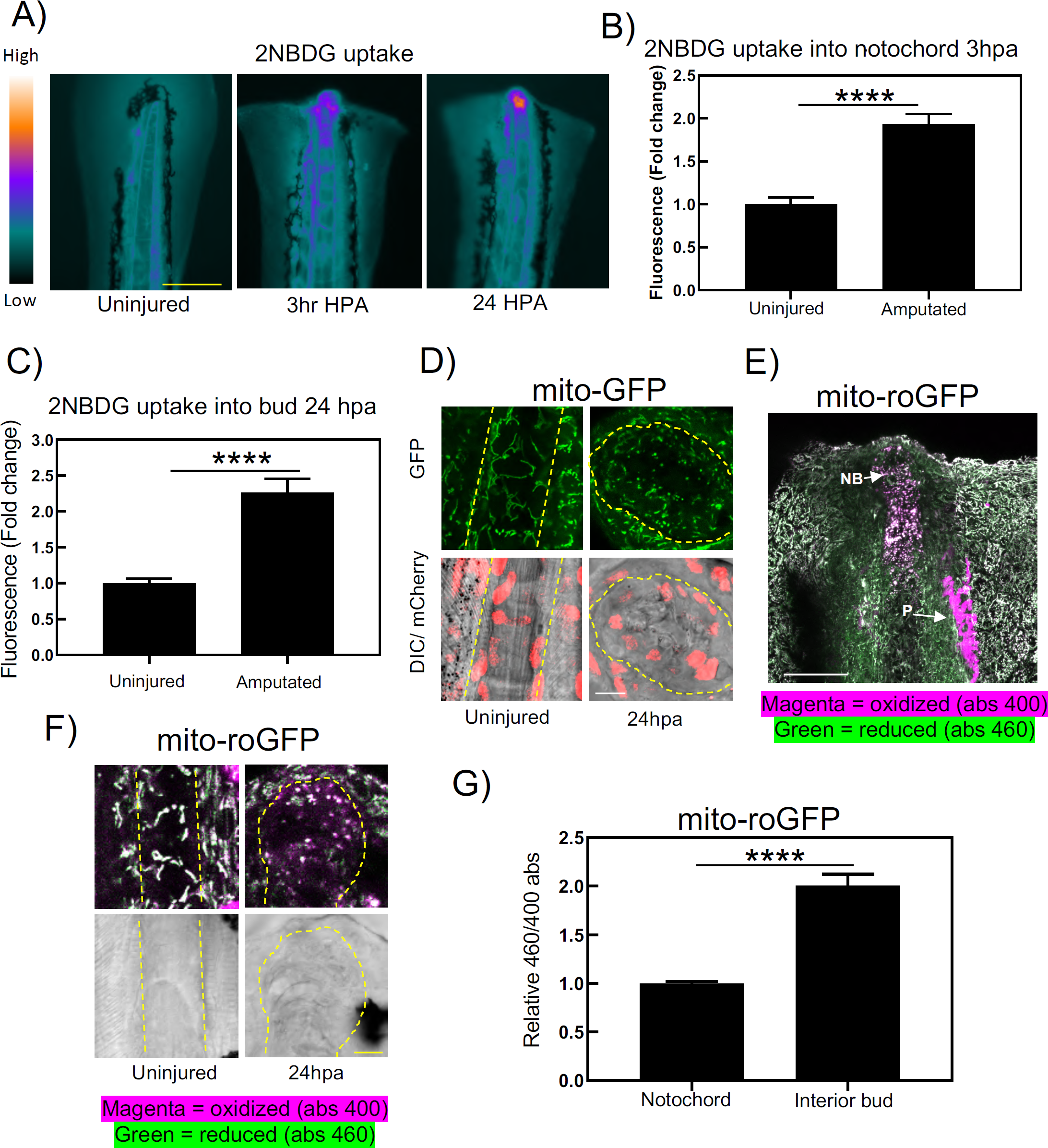
Notochord cells have altered metabolism during regeneration. A) Heatmap of fluorescent intensity of uninjured, 3 HPA, and 24 HPA embryos pulsed with 500 µM 2-NBDG for 2 hrs. Scale bar = 200 µM. B) Quantification of 2-NBDG fluorescence in notochord of uninjured and 3 HPA embryos. C) Quantification of 2-NBDG fluorescence in notochord of uninjured embryos and notochord bud of 24 HPA embryos. D) Confocal images of Tg(*actb2*:mito-GPF; *actb2*:H2a-tagRFP-T) embryos. Mitochondrial fragmentation can be seen in the cells comprising the blastema. Scale bar = 10 µM. E) Confocal image of Tg(*actb2*:mito-roGPF). NB indicates notochord bud. P indicates pigment autofluorescent pigment cells which are not depicting roGFP. A clear region of oxidized mitochondria can be seen in the notochord bud. Scale bar = 50 µM. F) Confocal images of roGPF in the notochord of uninjured embryos or blastema of 24 HPA embryos. The mitochondria remain highly oxidized. Scale bar = 10 µM. G) Quantification of relative absorbance of roGFP at 460 or 400 nm in the notochord of uninjured embryos or the notochord bud of 24 HPA embryos.

### scRNA-seq indicates that notochord cells transition to blastema through an EMT

Our data indicated that notochord cells undergo metabolic modifications consistent with glycolysis early in the regenerative process, and that glycolysis is essential for caudal fin regeneration, at least in part, through blastema formation. In order to understand the relationship between the notochord, the blastema, and how glycolysis is involved triggering regeneration, we performed scRNA-seq on 24 HPA and 48 HPA caudal fins with and without 2-DG treatment (Figure 4A, Supplementary Figure 2A). Based on gene expression in the untreated sample, we were able to identify specific populations of cells consistent with cell types expected to be present in the regenerating tail, including notochord and blastema cells (Figure 4B, Supplementary Figure 2B, Supplementary Table 1). A t-SNE analyses clustered blastema cells primarily into cluster 2, while notochord bud cells segregated into cluster 4 (Figure 4C). Several genes expressed in blastemas such as *aldh1a2, msx3*, and *msx1b* were also expressed in the mesenchyme. Additionally, other mesenchymal markers were also expressed in the blastema, supporting previous reports suggesting that the blastema is a mesenchymal tissue (Supplementary Figure 2C). Interestingly, there was a small group of cells which clustered mainly as blastema but overlapped with the notochord cell cluster (Figure 4D). While this population of cells expressed notochord markers, there were unexpectedly genes that were significantly enriched specifically within this small population (Figure 4E, Supplementary Table 2). The most significantly enriched gene was transgelin (*tagln*), a gene involved in motility which has been implicated in EMT and endothelial to mesenchyme transitions (endT). The most statistically overrepresented GO terms associated with genes expressed in this population were related to extracellular matrix organization, suggesting these cells may be gaining motility (Supplementary Table 3). Finally, these cells had a reduction in expression of notochord expressed genes and the epithelial maker desmoplakin (*dspa*), but an increase in expression of genes involved in EMT (Figure 4F). Altogether, these data indicate that the notochord contributed directly to blastema through an EMT and possible dedifferentiation.

**Figure 4:**
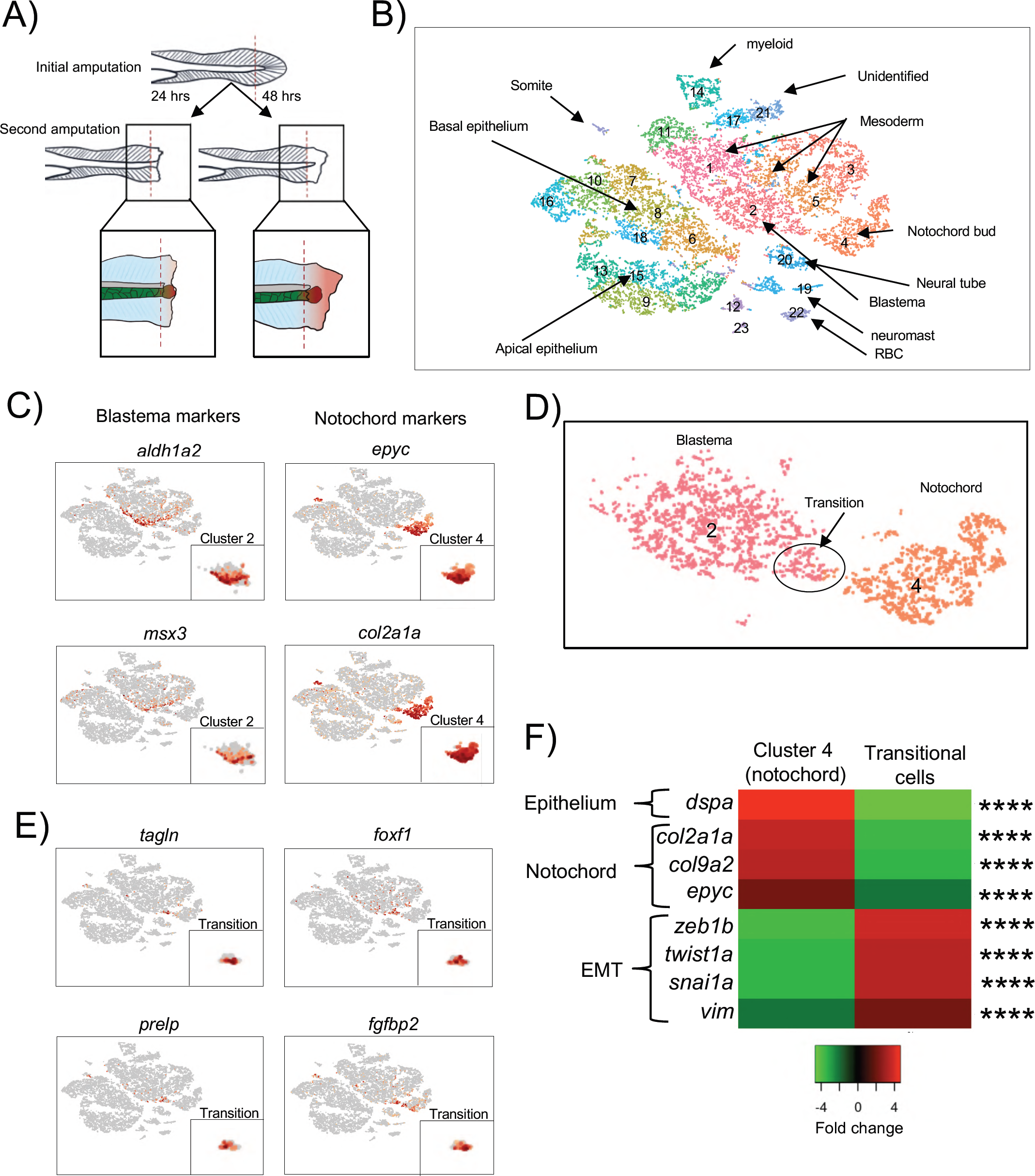
Blastema is formed from notochord cells. A) Illustration depicting the region of tail taken for scRNA-seq. B) t-SNE plot of cells from the scRNA-seq data. Number indicates cluster number. C) Gene expression of blastema markers and notochord markers in cells clustered by t-SNE. D) Portion of t-SNE plot depicting transitional population between notochord cells (cluster 4 from Figure 4B) and blastema cells (cluster 2 from Figure 4B). E) Gene expression of genes enriched in the notochord–blastema transitional population in cells clustered by t-SNE.

### Notochord sheath cells migrate out of the notochord bud to form the blastema

To further examine the relationship between notochord and blastema, we utilized a double transgenic zebrafish line, Tg(*col8a1a*:GFP; *col9a2*:mCherry) which expresses GFP in the vacuolated notochord cells and mCherry in the surrounding sheath cells [43]. Using time-lapse microscopy, we could show both cell types repositioned into the notochord bud post-amputation, generally maintaining the notochord architecture with vacuolated cells located on the interior of the bud surrounded by sheath cells (Figure 5A, Supplementary Movie 2). However, a time-lapse beginning at 24 HPA revealed sheath cells on the periphery of the notochord bud became motile and migrated out of the bud, suggesting these cells were undergoing a transformation to a different, more mobile cell type (Supplementary Movie 3, Supplementary Movie 4). Furthermore, these cells showed decreased and punctate (likely targeted for degradation) *col9a2*:mCherry expression, further indicating a loss of sheath cell identity (Figures 5B and 5C). By 48 HPA these transformed cells had migrated throughout the regenerating fin (Figure 5D). Based on this evidence, we hypothesized that the sheath cells on the periphery of the notochord bud were the transitional cells between notochord and blastema in our scRNA-seq data. To further evaluate, we analyzed double transgenic zebrafish expressing *col9a2*:mCherry and *tagln*:GFP [44]. In uninjured animals, no GFP is detectable in sheath cells. However, at 24 HPA we observed strong *tagln*:GFP expression specifically in sheath cells that also had the reduced, punctate mCherry expression on the periphery of the notochord bud (Figure 5E, 5F). Together, our data show notochord sheath cells undergo dedifferentiation and an EMT to become mesenchymal blastema cells.

**Figure 5:**
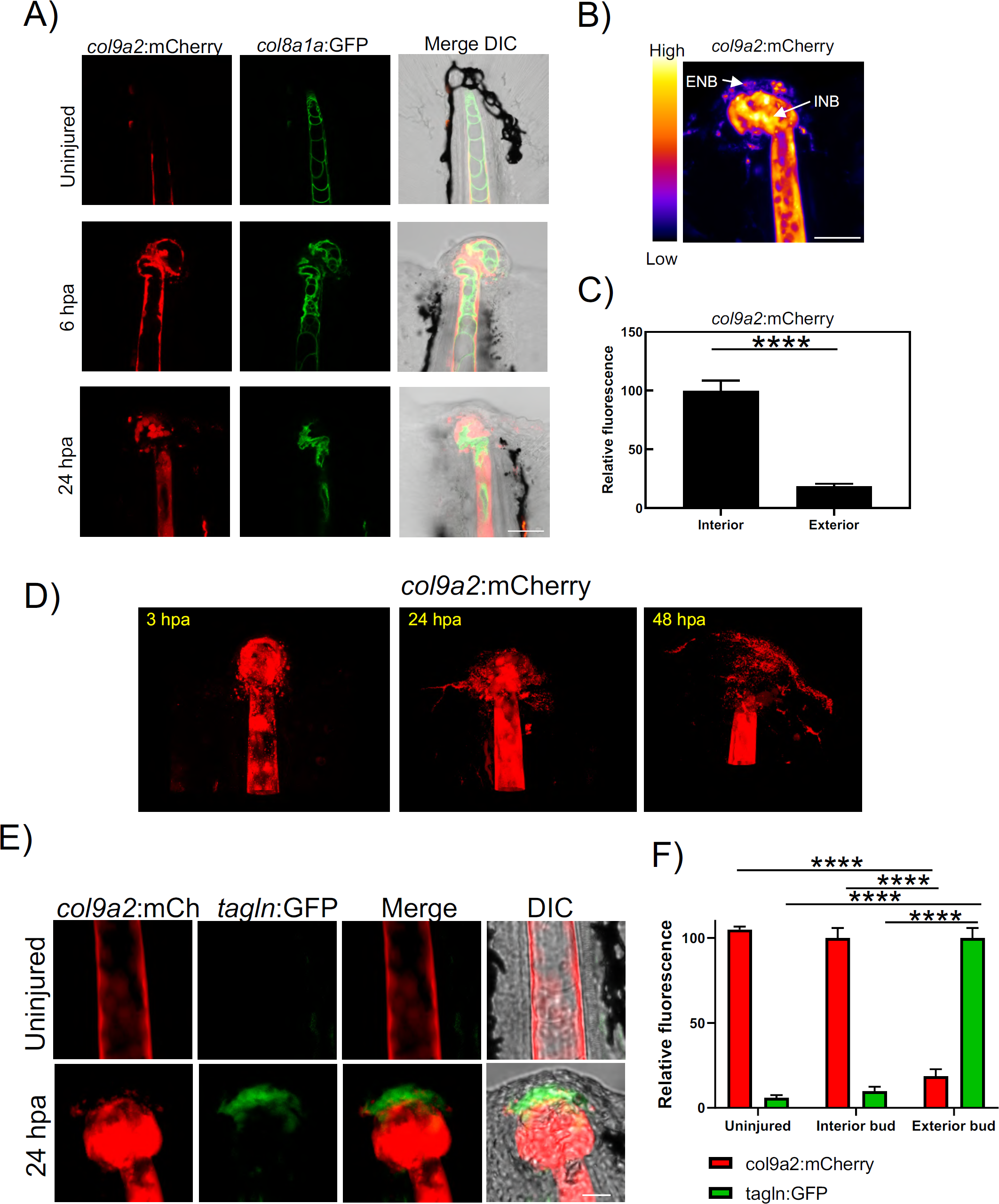
Notochord sheath cells form blastema through an EMT. A) Confocal images depicting tails of uninjured, 6 HPA, or 24 HPA Tg(*col9a2*:mCherry; *col8a1a*:GFP) embryos. Scale bar = 50 µM. Green are notochord cells, red are sheath cells. B) Heatmap of mCherry (sheath cell) fluorescence in tail of 24 HPA Tg(*col9a2*:mCherry) embryo. Scale bar = 50 µM. INB indicates interior notochord bud, ENB indicates exterior notochord bud. Cells on the periphery are lower in red fluorescence. C) Quantification of mCherry fluorescence in interior and exterior of notochord bud of 24 HPA Tg(*col9a2*:mCherry) embryo. D) Images depicting migration of sheath cells out of the notochord bud from 3 to 48 HPA. Sheath cells are expressing *col9a2*:mCherry. E) Confocal images depicting the notochord of uninjured or the notochord bud of 24 HPA Tg(*col9a2*:mCherry; *tagln*:GFP) embryos. Scale bar = 20 µM. Transgelin positive cells are detectable in the weaker red fluorescent cells on the periphery. F) Quantification of GFP and mCherry fluorescence in Tg(*col9a2*:mCherry; *tagln*:GFP) embryos. Fluorescence in the notochord of uninjured embryos and the exterior and interior notochord bud of 24 HPA embryos was measured.

### TGF–β signaling is required for blastema formation

We next sought to understand how glycolysis regulates blastema formation. Visualizing our scRNA-seq as a t-SNE plot by library showed that control and 2-DG treated cells formed distinguishable groups within the blastema cluster, indicating that 2-DG strongly altered gene expression of some genes in the blastema primarily preventing genes from being expressed (Figure 6A, Supplementary Table 4). Among the genes suppressed by 2-DG were blastema markers *aldh1a2* and *msx3*, confirming our previous data suggesting that the blastema does not form properly when glycolysis is inhibited (Figure 6B). In order to understand how inhibition of glycolysis results in aberrant blastema formation, we examined pathways that were significantly overrepresented among genes suppressed by 2-DG in the blastema. As expected because of the reduced cell division in 2-DG treated cells (Figure 2E,F), genes in the de novo pyrimidine deoxyribonucleotide biosynthesis pathway, which is important for DNA replication and therefore cell proliferation, are overrepresented (Figure 6C). However, genes involved in TGF–β signaling, a well-characterized inducer of EMT, were also significantly overrepresented. Interestingly, components of TGF–β signaling were strongly expressed in the notochord bud (cluster 4), indicating TGF–β signaling may be required for sheath cell EMT prior to blastema formation (Figure 6D). We were able to show that inhibition of TGF–β signaling blocked regeneration beyond formation of the notochord bud, and similar to 2-DG, resulted in repression of *msx3* (Figures 6E, 6F, 6G). Interestingly, unlike 2-DG treatment, cell proliferation continued in the presence of the TGF–β inhibitor, suggesting TGF–β signaling is necessary for sheath cell EMT but the signal is downstream of both the glycolytic shift and the initiation of cell division (Figure 6H and 6I).

**Figure 6:**
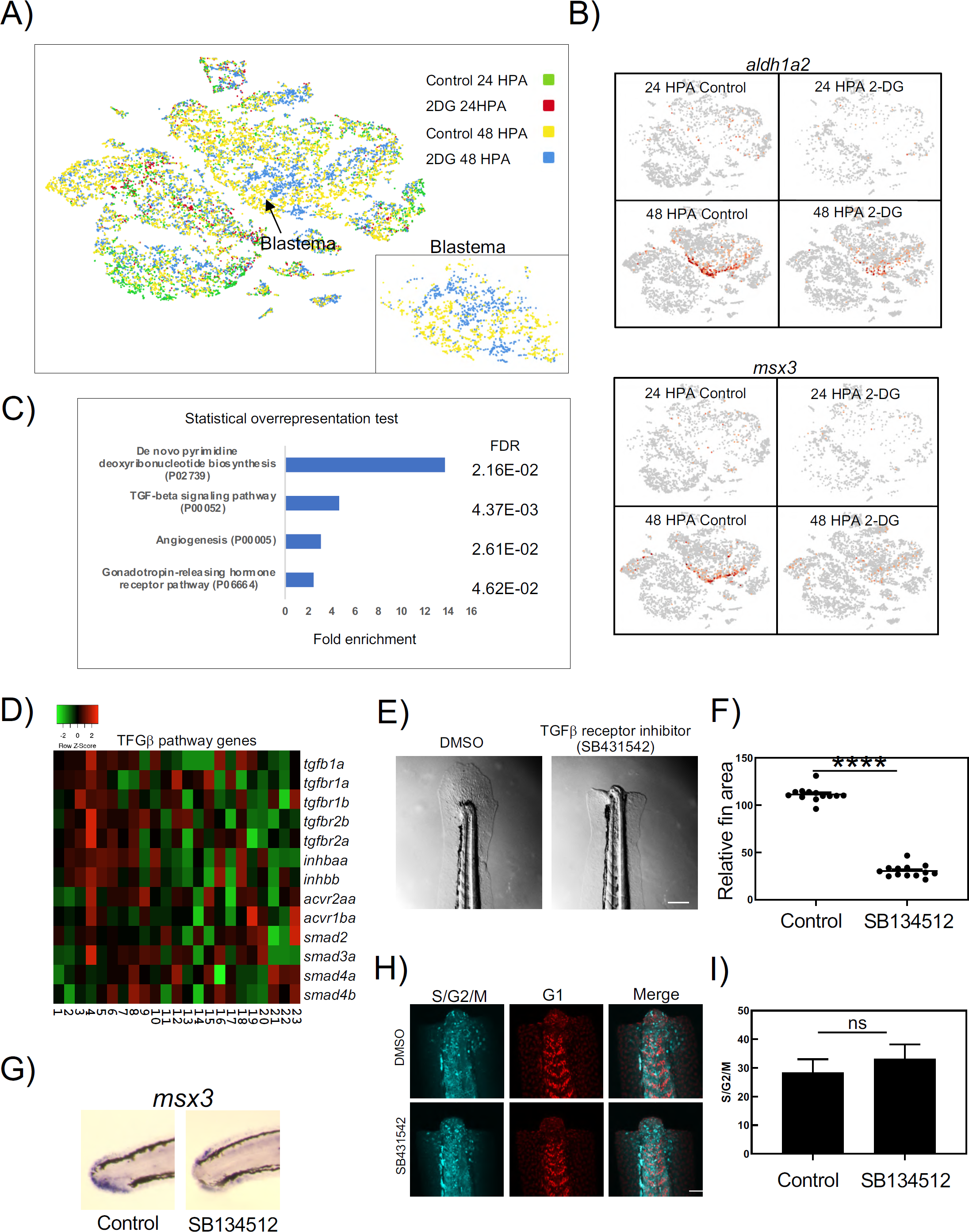
TGF–β signaling is essential for development of blastema. A) t-SNE plot of cells from scRNA-seq shown by library. B) Expression of blastema markers *aldh1a2* and *msx3* shown on a t-SNE in each library. Both genes were suppressed by 2-DG in the blastema, p<0.001. D) Relative expression of genes involved in TGF–β signaling by clusters generated by t-SNE. E) Image of the tail of a 96 HPA embryo treated with DMSO or the alk4, 5, and 7 inhibitor SB431542. Drug was added to embryo media 2 hours prior to amputations and treatment persisted through the duration of experiment. Scale bar = 200 µM. F) Quantification of tail area of 96 HPA embryos treated with DMSO or SB431542. Drug was added to embryo media 2 hours prior to amputations and treatment persisted throughout the duration of experiment. G) In situ hybridization of *msx3* in DMSO or SB431542 treated embryos. H) Confocal images of DMSO or SB431542 treated Fucci fish embryo tails at 24 HPA. Scale bar = 50 µM. I) Quantification of cells in S, G2, or M phase in DMSO or SB431542 treated Fucci embryo tails at 24 HPA.

### A metabolic shift is necessary to activate a TGF–β induced EMT

Our data indicated that EMT was necessary for blastema development and that a glycolytic shift is necessary to induce TGF–β signaling, a major driver of EMT. Specifically, expression of two activin subunits, *inhbaa* and *inhbb*, were repressed by 2-DG in the notochord bud at 24 hpa, while *inhbaa, inhbb, tgfb1*, and *tgfbr2b* were repressed in the blastema at 48 hpa (Figure 7A). Additionally, both zebrafish paralogs of *snai1*, a mediator of TGF–β induced EMT, were suppressed by 2-DG in the blastema. To examine whether the transcriptional suppression of TGF–β genes by 2-DG reflected an actual reduction in protein signaling, we performed immunofluorescence using antibodies to phospho-SMAD2 (P-SMAD2). In support of our scRNA-seq data, caudal fin amputation resulted in strong activation of TGF–β signaling in the notochord bud by 24 HPA, followed by signaling in the blastema at 48 HPA (Figure 7B). As expected, activation was reduced with the TGF–β inhibitor SB431542, indicating the P-SMAD2 antibody staining was specific for TGF–β signaling (Supplementary Figure 3). At 48 HPA, we observed a decrease in nuclear P-SMAD2 staining in the blastema upon 2-DG treatment, consistent with repression of TGF–βeta ligands expressed in the blastema (*inhbaa, inhbb, tgfb1*) (Figure 7C). We then analyzed the *snai1a* expression to probe whether TGF–β signaling was required for EMT during tail regeneration. Both 2-DG and SB431542 repressed *snai1a* expression, suggesting a block in EMT with either treatment (Figure 7D). Together, our data indicate that the glycolytic shift was necessary to induce TGF–β signaling, which promotes blastema formation through dedifferentiation and EMT of notochord sheath cells (Figure 7E)

**Figure 7:**
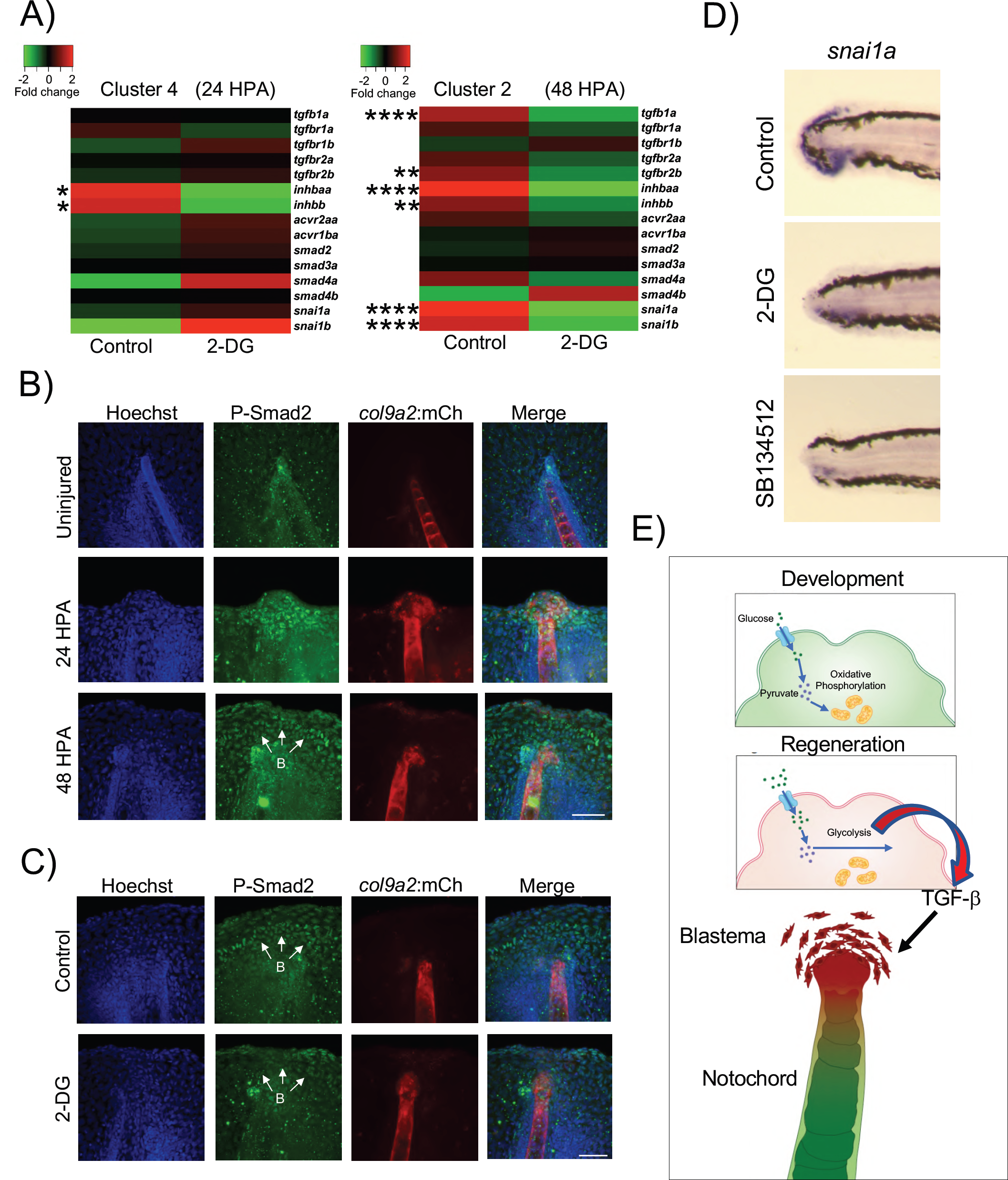
Glycolysis promotes TGF–β signaling in blastema. A) Heat maps of gene expression in scRNA-seq clusters 2 (blastema) and 4 (notochord bud). ****P < 0.0001, ***P < 0.001, **P < 0.01, *P < 0.05. B) Immunofluorescent staining of P-SMAD2 in uninjured and 24 HPA and 48 HPA Tg(*col9a2*:mCherry) embryos. Scale bar = 50 µM. C) Immunofluorescent staining of P-SMAD2 in 48 HPA Tg(*col9a2*:mCherry) embryos with (control) or without 2-DG treatment. B indicates blastema. Scale bar = 50 µM. D) In situ hybridization of *snail1a* in 48 HPA embryos. E) Model of blastema formation during tail regeneration in zebrafish embryos. A metabolic shift to glycolysis promotes TGF–β signaling, which results in EMT of notochord sheath cells to form the blastema.

## Discussion

Despite work done to show that blastema arises from dedifferentiation of existing “fate committed” cells[45, 46], how this process occurs has remained largely unknown. We have shown that in the embryonic zebrafish tail, normal blastema formation requires a metabolic shift to glycolysis. This shift promotes TGF–β signaling, resulting in blastema formation through dedifferentiation and an EMT of notochord sheath cells, which go on to regenerate all the various cell types in the tail fin-fold.

The Warburg effect was initially thought of as a process unique to cancer metabolism, however, it has become clear that metabolic switching has key regulatory roles under many normal physiologic conditions [47]. It has been assumed that highly proliferative cells require fast ATP production and have an increased need for macromolecule synthesis, but there is also evidence that glycolysis is utilized for cell differentiation or dedifferentiation depending on biological context [48, 49]. Our data indicate that in the regenerating zebrafish tail, glycolysis is required for both cell proliferation and to trigger dedifferentiation through TGF–β signaling. Proliferation and dedifferentiation are not necessarily mutually exclusive processes and we see evidence that the dedifferentiation can be decoupled from cell proliferation. We observe a decrease in expression of several regulators of cell proliferation in the blastema upon 2-DG treatment and hypothesize that the blastema is not forming properly and therefore is not competent for increased proliferation. This is supported by our observation that other tissues such as the neural tube, which do not undergo a metabolic shift, still express markers of proliferating cells during regeneration despite the 2-DG treatment. Therefore, glycolysis-dependent proliferation may not be a generalized phenomenon but reliant on specific signaling pathways in the context of blastema formation.

The activin genes *inhbaa* and *inhbb* are initially transcriptionally repressed by 2-DG at 24 HPA in the notochord bud, but *tgfb1* is not repressed until later in the regenerative process. Accordingly, by assaying phospho-SMAD2 activity, we did not observe a decrease in overall TGF–β signaling with 2-DG treatment until 48 HPA in the blastema, when both the activin genes and *tgfb1* were repressed. Interestingly, although not statistically significant due to small size of the population, *tgfb1* also appears to be repressed by 2-DG treatment in the transitional population between notochord cells and blastema. Furthermore, *inhbaa* expression, which was dependent on the metabolic shift to glycolysis, was highest in these transitional cells when compared to either the notochord bud or the blastema. Therefore, this is likely the key population where glycolysis strongly promotes TGF–β signaling. This potentially explains why an aberrant, non-proliferative blastema, rather than no blastema, forms when glycolysis is inhibited. We hypothesize that initial TGF–β signaling through TGFB1 begins the dedifferentiation process and is independent of glycolysis, but the glycolytic shift is necessary to maintain a sustained TGF–β signal via activin which initiates the EMT, promotes cell proliferation, and makes the blastema competent for coordinated regeneration. In support of this hypothesis *snai1a*, which is upregulated in these transitional cells compared to the notochord bud and remains strongly expressed in the blastema, is suppressed by both inhibition of glycolysis or inhibition of TGF–β signaling.

A previous report demonstrated that notochord streaming was triggered through ROS driven hedgehog signaling and was necessary for regeneration [21]. However, we have shown that the contribution of the notochord to regeneration goes beyond being solely a source for signaling ligands, as sheath cells directly transition to blastema cells in a TGF–β dependent process. ROS generation and glycolysis are both important contributors to cellular redox state, and reflect the complex, interdependent nature of cellular responses to injury. ROS and glycolysis have been shown to reciprocally regulate each other, and both pathways are required for functional blastema formation [21, 50]. However, our data, in contrast to ROS, show no evidence that glycolysis regulates sonic hedgehog ligands in the notochord bud or blastema. This suggests that ROS and glycolysis are activating distinct, parallel pathways, or alternatively ROS and sonic hedgehog acts upstream of the glycolytic shift and the subsequent TGF–β signaling. Because 2-DG inhibits glycolysis at the very first step (glucose-6-phosphate), further research will be required to tease apart whether it is glycolysis, the related pentose phosphate pathway (PPP), or the hexosamine biosynthetic pathway that provides the key cellular signal for sustaining TGF–β signaling in the blastema (Supplementary Figure 4).

Sheath cells being the primary contributing cell type of a forming blastema is clearly limited to embryonic tail regeneration where a notochord is still present. The source of blastema cells must certainly differ depending on the species, the tissue, and age of the animal, but there are likely conserved mechanisms controlling dedifferentiation of surrounding cells and blastema formation. Glycolysis and the PPP have been proposed to be preferred to oxidative phosphorylation during wound healing and early blastema phases during both lizard and xenopus tail regeneration [5, 6]. Additionally, TGF–β is required for EMT and blastema formation in lizard tail regeneration, zebrafish fin regeneration [51], and axolotl limb regeneration [52, 53]. Interestingly, in adult zebrafish and gecko tails, activin appears to be the primary TGF–β ligand, while in our dataset, both activin and TGF-β1 are inhibited by 2-DG treatment [51, 52]. It is possible that activin induced by metabolic shift during regeneration is a conserved mechanism in regenerative species.

With respect to mammals, the MRL strain of mouse, which has superior regenerative capacity compared to wild-type mice, has higher levels of aerobic glycolysis than normal mice, potentially preserving some modest ability to trigger blastema formation, at least under limited circumstances [54]. Additionally, several studies indicate that a glycolytic shift is required for dedifferentiation of mammalian cells in vitro [49, 55, 56]. Therefore, it is likely that our findings will contribute to our understanding of regeneration and wound healing on a broader scale.

In addition to tissue regeneration and stem cell biology, blastema formation shows several parallels to cancer biology, including a shift to glucose-dependent metabolism and an induction of an EMT followed by cell migration. However, unlike tumor progression, regeneration is spatially and temporally coordinated, allowing us to control variables and obtain insight into critical but transient gene expression within sub-populations of cells. By using scRNA-seq, we were able to capture notochord sheath cells in a transitional state on their way to becoming the mesenchymal blastema. The most significantly enriched gene in this population was transgelin (*tagln*). Transgelin is only transiently expressed during the EMT, whereas many of the main drivers of EMT such as *snai1* continued to be expressed in the blastema. Transgelin is an actin binding protein expressed in smooth muscle, fibroblasts, and the umbilical cord mesenchymal stem cells. Furthermore, its expression has been detected in cancer stem cells and has been implicated in their migration and invasion [57]. Paradoxically, trangelin has been described as both promoting and inhibiting bladder cancer [58, 59]. The paradox is potentially resolved if timing and transient transgelin expression is important in the progression of bladder cancer as it is in the formation of a blastema. Due to many observed similarities between wound healing and cancer, cancer has been described as a wound that does not heal [60]. Moreover, multiple studies showing that both wound healing (or regeneration) and cancer can have similar transcriptional profiles [61, 62]. Therefore this work may provide insight into the cellular processes that drive tumor progression as well as tissue regeneration [6, 52, 53, 63].

## Acknowledgements

We thank M. Bagnat (Duke University) for sharing the Tg(*col8a1a*:GFP; *col9a2*:mCherry) zebrafish line, D. Kimelman (University of Washington) for the dual Fucci zebrafish line, and M. Pack (University of Pennsylvania) for sharing the Tg(*tagln*:GFP) zebrafish line. The roGFP entry clone was a kind git from K. Drerup (National Institute of Child Health and Development). Jennifer Wang provided illustrations for this manuscript. We have complied with all relevant ethical regulations regarding animal use and all animal experiments were approved by the National Human Genome Research Institute’s Animal Care and Use Committee (protocol #G-01-3). This research was supported by the Intramural Research Program of the National Human Genome Research Institute (ZIA HG200386-08).

## Figure Legends

**Supplementary Figure 1:**
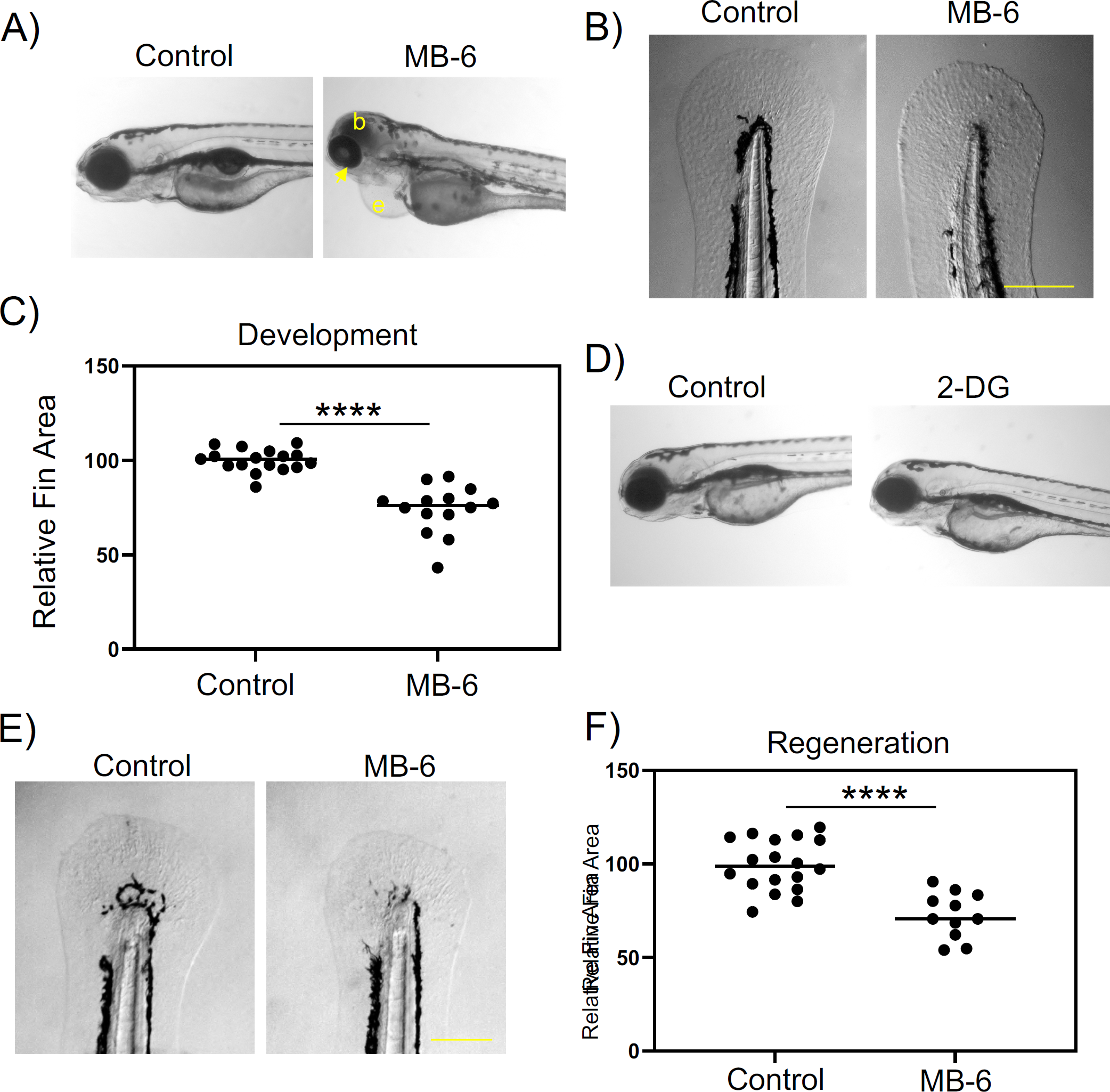
Mitochondrial function is essential for embryonic development. A) Images of 4 DPF embryos treated with 1% DMSO or 2.5 µM MB-6 from 1-4 DPF. b = blood pooling in the head. e= edema. Arrow indicates underdeveloped eye. B) Images of tails of 4 DPF embryos treated with 1% DMSO or 2.5 µM MB-6 from 1-4 DPF. Only minor differences in size could be detected. Embryos were treated 1 DPF through the duration of the experiment. Scale bar = 200 µM. C) Quantification of tail surface area at 4 DPF of embryos treated with 1% DMSO or 2.5 µM MB-6. D) Images of 4 DPF embryos untreated (control) or treated with 100 mM 2-DG from 1-4 DPF. E) Images of tails of 7 DPF embryos, 4 days post-amputation treated with 1% DMSO or 2.5 µM MB-6. F) Quantification of tail surface area at 96 HPA of embryos treated with 1% DMSO or 2.5 2.5 µM MB-6. Embryos were treated 2 hours before amputation through the duration of the experiment.

**Supplementary Figure 2:**
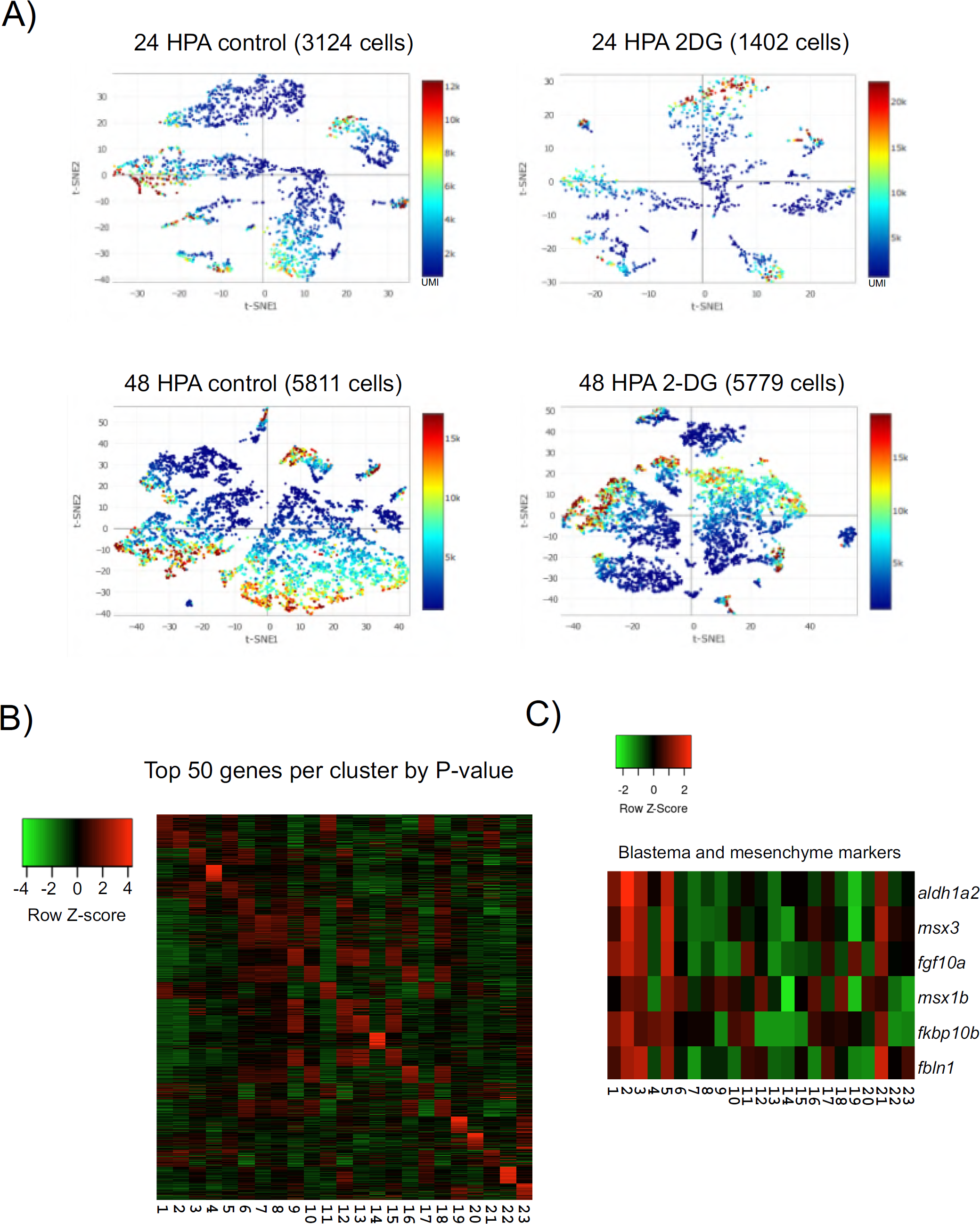
scRNAseq of regenerating tail. A) t-SNE plots from each library color coded by unique molecular identifier (UMI) counts. B) Heatmap of relative gene expression of the top 50 markers for each cluster as determined by p-value. C) Heatmap of relative gene expression for epithelial, notochord, and EMT markers in notochord cells or notochord− blastema transitional cells. D) Heatmap of gene expression of blastema of mesenchyme markers by t-SNE generated clusters. Columns depict cluster number.

**Supplementary Figure 3:**
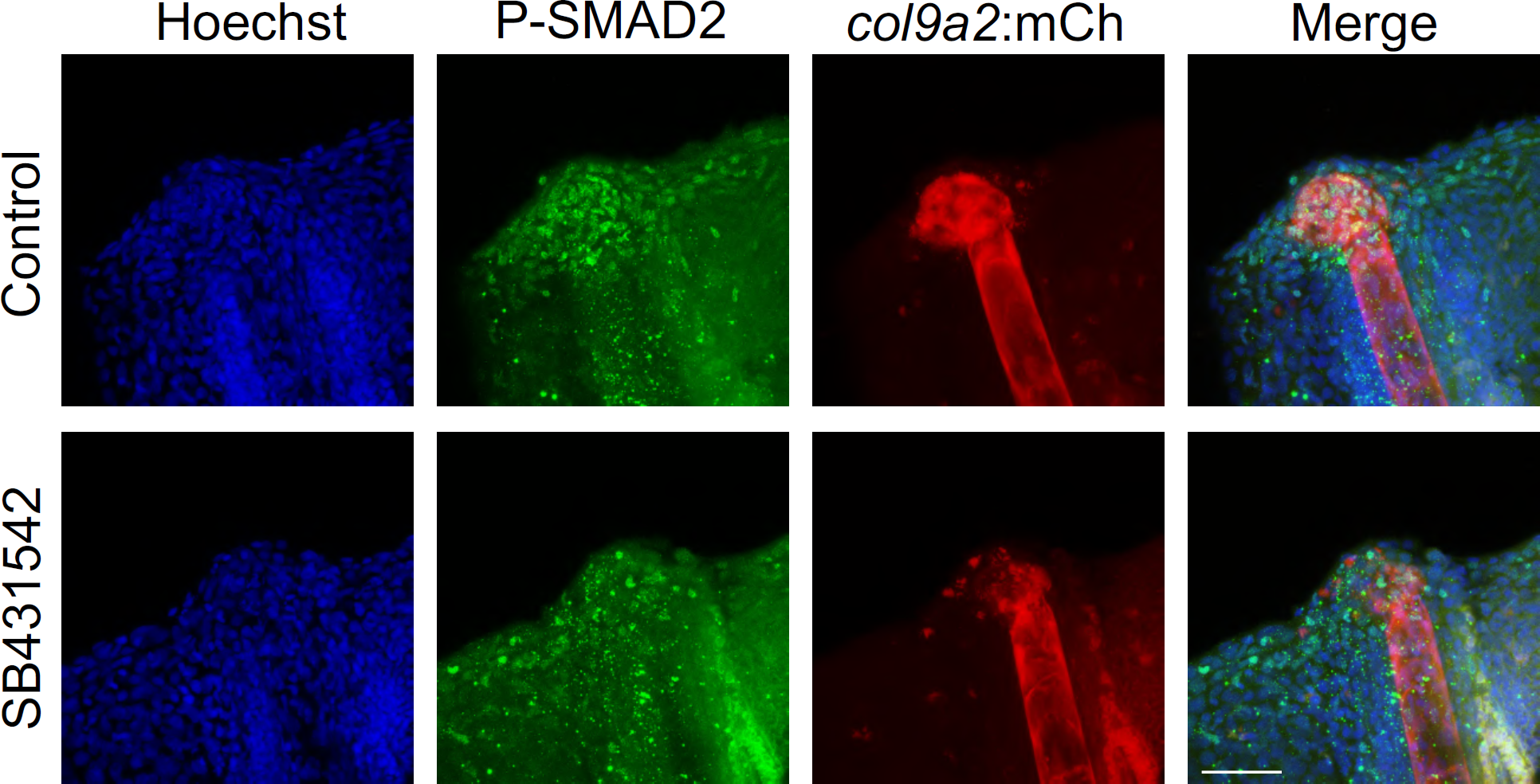
SB431542 reduces SMAD2 phosphorylation. Immunofluorescent staining of P-SMAD2 in uninjured and 24 HPA Tg(*col9a2*:mCherry) embryos. Scale bar = 50 µM.

**Supplementary Figure 4:**
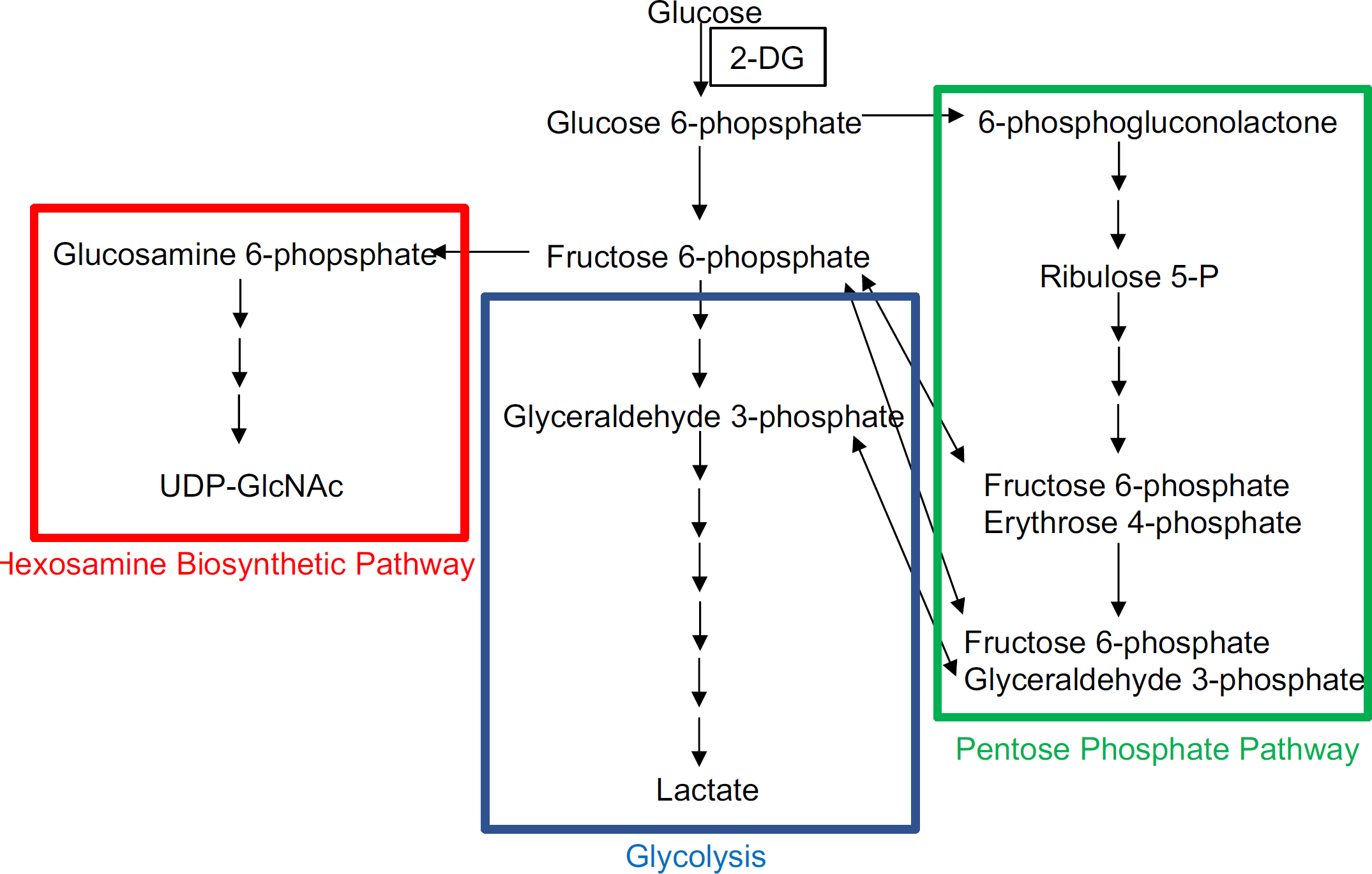
Glycolysis and related pathways. Diagram depicting glycolysis, the pentose phosphate pathway, and the hexosamine biosynthetic pathway. Single-head Arrows indicate pathway intermediates / products. Double-headed arrows indicate shared pathway substrates.

